# Robust spatial self-organization in crowds of asynchronous pedestrians

**DOI:** 10.1101/2023.08.02.551584

**Authors:** Takenori Tomaru, Yuta Nishiyama, Claudio Feliciani, Hisashi Murakami

## Abstract

Similar to other animal groups, human crowds exhibit a variety of self-organized collective behaviors. Spontaneous formation of unidirectional lanes in bidirectional pedestrian flows is one of the most striking examples of self-organization in human crowds. In addition, previous experimental studies have suggested that stepping among pedestrians is spontaneously synchronized over time. However, those studies have primarily focused on one-dimensional, single-file crowds, limiting our understanding of temporal stepping patterns in crowds with more freedom of movement and their relationship to spatial patterns of organization. Here, we conducted experiments of bidirectional pedestrian flows (24 pedestrians in each group) and investigated the relationship between collective footsteps and self-organized lane formation by tracking each individual trajectory as well as foot-stepping kinematics. We found that, unlike previous studies, pedestrians did not spontaneously align their steps unless there was an external auditory cue to follow. Moreover, we found that footstep synchronization generated by external cues disturbed the flexibility of pedestrians’ lateral movements, increased potential collision risks, and narrowed the formed lanes, suggesting structural instability in the spatial organization. These results imply that, without external cues, pedestrians marching out of step with each other can efficiently self-organize into robust structures. Understanding how asynchronous individuals contribute to ordered collective behavior has potential applications for promoting efficient crowd management and will inform theoretical models for human and other animal collective behavior with asynchronous positional updates.

## Introduction

Highly organized collective behavior exhibited by a large number of people, as seen in a marching band, impresses spectators. Such disciplined behavior is orchestrated through a predetermined plan and/or guided by an external conductor from a global perspective. However, collective behavior can also organize spontaneously in nature. Witnessing massive flocks of birds and schools of fish can be an amazing sight [1]. It is also fascinating that many people can come and go on crowded streets of cities worldwide without colliding. In these behaviors, there are no predetermined plans or external conductors, and individuals behave relatively freely. Instead of being driven by external global forces, they self-organize through local interactions among group members [2].

Human crowds, in particular, have attracted attention from researchers in various fields, with the aim of helping to manage mass events and daily pedestrian transportation [3]. Some collective patterns of organization bring about beneficial results to the group, such as lane formation, where unidirectional lanes are spontaneously formed in bidirectional pedestrian flows in crowded streets or crossings, which increases the efficiency of traffic flow [4–7]. On the other hand, if people in a crowd self-organize in other ways, such as crowd turbulence, they become uncontrollable and can lead to serious disasters [8, 9]. Although providing traffic information and guidance is important to proactively prevent crowd disasters, it is difficult to restrict and control the movements of people who have happened to gather together [3]. To accomplish crowd management that facilitates traits of pedestrian’s movements that contribute to functional self-organization, it is essential to understand the mechanisms underlying collective human behavior.

Some recent experiments have suggested that the footsteps of pedestrians in a group are synchronized. For example, their footsteps can spontaneously align even without indirect interactions among pedestrians via a wobbling walkway [10–14], unlike the case of London Millennium Bridge, where pedestrians fell into step via vibrations of the suspension footbridge [15–17]. Most previous studies on spontaneous footstep synchronization in human crowds have adopted a one-dimensional approach known as the single-file experiment [12–14]. In this setup, pedestrians walk in the same direction along a narrow, circular-shaped corridor, which prohibits lateral movements and overtaking and enables researchers to extract elementary forms of interactions between consecutive pedestrians. The footstep synchronization observed in this manner can serve as an optimization strategy, where simultaneous movements of each pedestrian’s same-side feet efficiently exploit limited spatial resources in a jammed situation, reducing collisions and enhancing overall flow [12–14]. The assumption of functional footstep synchronization has also been supported by various single-file experiments where pedestrians walk while listening to or following external sounds (e.g., music or a sound at a constant tempo) [18–20].

However, in the one-dimensional approach, the stability of the single-file structure is guaranteed by a boundary condition (i.e., a narrow corridor with borders). In contrast, most pedestrians’ motions in daily life are not restricted to one dimension, and pedestrians can spontaneously generate ordered structures in the absence of external forces, such as those observed with lane formation phenomena [4–7]. In addition to making adjustments to preceding and following individuals, pedestrians commonly use lateral movements under two-dimensional conditions [4, 6]. Pedestrians dynamically modify their configurations to avoid collisions with oncoming pedestrians and to overtake others. It is possible that footstep synchronization may be an optimal strategy to exploit spatial resources with a preceding individual only when the single-file structure remains static. To further investigate the role of collective footsteps in human crowds, it is essential to investigate pedestrian behavior under conditions that allow them to move with more freedom of movement than available in one-dimensional experiments.

In this study, we conducted experiments to examine lane formation phenomena in a crossing scenario by means of tracking pedestrians’ positions in two dimensions and recording their foot movements. Lane formation can be an ideal system to investigate footstep synchronization in two dimensions because pedestrians spontaneously segregate into multiple lanes, which are not fixed static structures, and lateral movements dynamically contribute to self-organization. In fact, previous experiments revealed the intrinsic instability of bidirectional flow organization due to overtaking behavior and lateral movements [4]. Furthermore, in another experiment, during the development of lanes, lateral explorations from the direct straight path to the destination inevitably occurred as pedestrians passed through a crowd, avoided oncoming pedestrians, and thereby achieved lane formation [6]. In these previous experiments, pedestrians did not follow any external cues to temporally adjust their movements, and moving at their own timing may have facilitated such lateral exploration. In other words, aligning steps with external cues might restrict lateral fluctuations in a two-dimensional scenario. To address this possibility, we set an experimental condition in which participants were asked to align their steps with external auditory cues and compared the results with those of a baseline condition without any temporal cues.

We find that pedestrians do not spontaneously align their footsteps with each other unless there is an external conductor. Moreover, although we observed that the external cues could produce an organized structure, they also yield potential instability, which was characterized by an increased number of pedestrian lanes formed and a shorter time/distance to potential collisions. Furthermore, we find that aligning steps with external cues decreases the lateral exploratory motion of pedestrians, suggesting a relation to the width of generated lanes. These findings shed light on the importance of asynchronous motions enabling exploratory lateral behavior and its contributions to robustly self-organizing human crowds; hence, they should inform theoretical models with asynchronous positional updates.

## Results

We conducted crowd experiments in which two groups (24 pedestrians each) bidirectionally walked in a mock corridor (Figs. 1A and S1, see Methods). In order to investigate the effect of external auditory cues on individual behavior and on self-organization of bidirectional flow, we set two experimental conditions: baseline (BASE) and MARCHING. In the MARCHING condition, all pedestrians were asked to walk to an electric metronome with a tempo of 120 beats/min (approximately equal to the normal pedestrian step frequency [19]) played from a loudspeaker. All pedestrians in one of the two groups were equipped two inertial measurement units (IMUs), one on each leg, in each condition. The experiments were replicated 20 times under each condition.

**Fig. 1.**
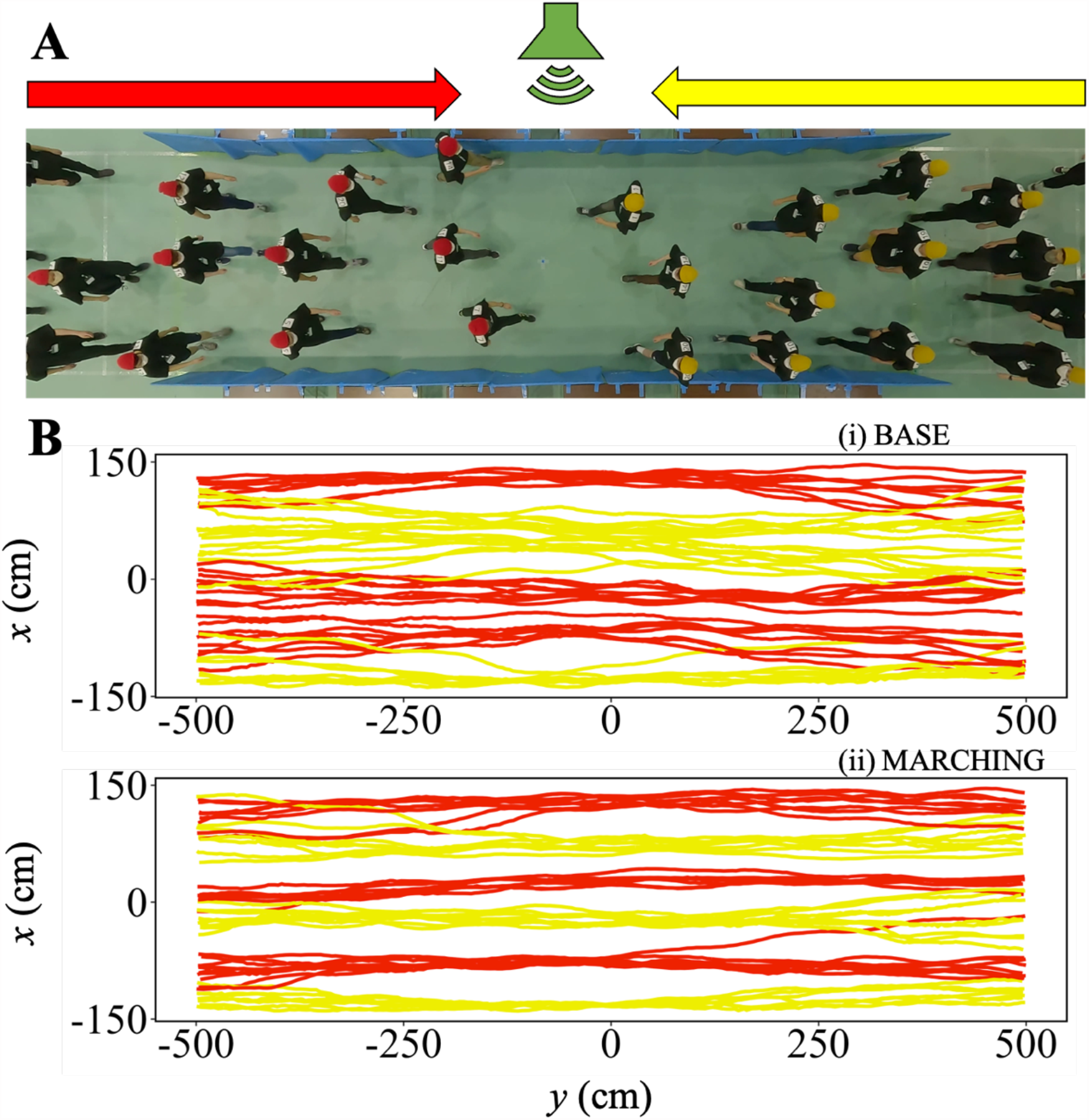
Bidirectional flow experiments with an external auditory cue. (A) Snapshot from an experiment. Auditory stimulus was played by the loudspeaker placed on one side of the experimental corridor under the MARCHING condition. (B) Examples of reconstructed pedestrian trajectories under the (i) BASE and (ii) MARCHING conditions. Yellow (red) lines represent pedestrians moving from right (left) to left (right).

Figure 1B shows representative examples of the reconstructed pedestrian trajectories under each condition. According to our observations, pedestrians in the two groups started to walk in opposite directions (toward each other) along the corridor and deviated from a direct straight path to their destinations as they sought passage through the crowd and to avoid oncoming pedestrians. In doing so, they self-organized into several unidirectional lanes. During experiments, footsteps of pedestrians under the MARCHING condition were apparently synchronized whereas those under BASE were not. Moreover, pedestrians under the BASE condition seemed to deviate more laterally and merge into fewer lanes (i.e., wider lanes) than those under the MARCHING condition.

### Influence of external auditory cues

To quantitatively verify the above observations, we estimated the timing of every heel strike from the IMUs and determined step synchronizations between pairs of two pedestrians for each trial (see [10, 12–14] and Methods). We identified the pairs so that pedestrian *i* was paired with pedestrian *j* if *j* was the individual that *i* followed for the longest time during a trial (see [4] for a more detailed definition of *following*). Figure 2A shows the proportion of synchronized footsteps out of all the paired footsteps for each trial. We found that the proportion of synchronized steps in the MARCHING condition was considerably higher than that in the BASE condition. To determine the extent to which they were synchronized more than by chance, especially in the BASE condition, we performed 1000 virtual trials for each condition, in which two pedestrians in pairs were randomly shuffled (i.e., a follower from one pair was paired with a preceding individual from another pair in the same condition). We then calculated the proportion of synchronized steps for this group and compared it to that of the original dataset (see Methods). There was a significant difference in the degrees of synchronization between original and random datasets in MARCHING (Fig. 2A, Welch’s *t*-test, *N*_*actual*_ = 20, *N*_*random*_ = 1000, *t* = 13.7, *p* < 0.001, Cohen’s *d* = 2.89), but not in BASE (Welch’s *t*-test, *N*_*actual*_ = 20, *N*_*random*_ = 1000, *t* = −1.08, *p* = 0.29, Cohen’ *d* = −0.23). This suggests that pedestrians spontaneously synchronize their footsteps with each other in the presence of an external cue, while they do not do so if the absence of an external cue.

**Fig. 2.**
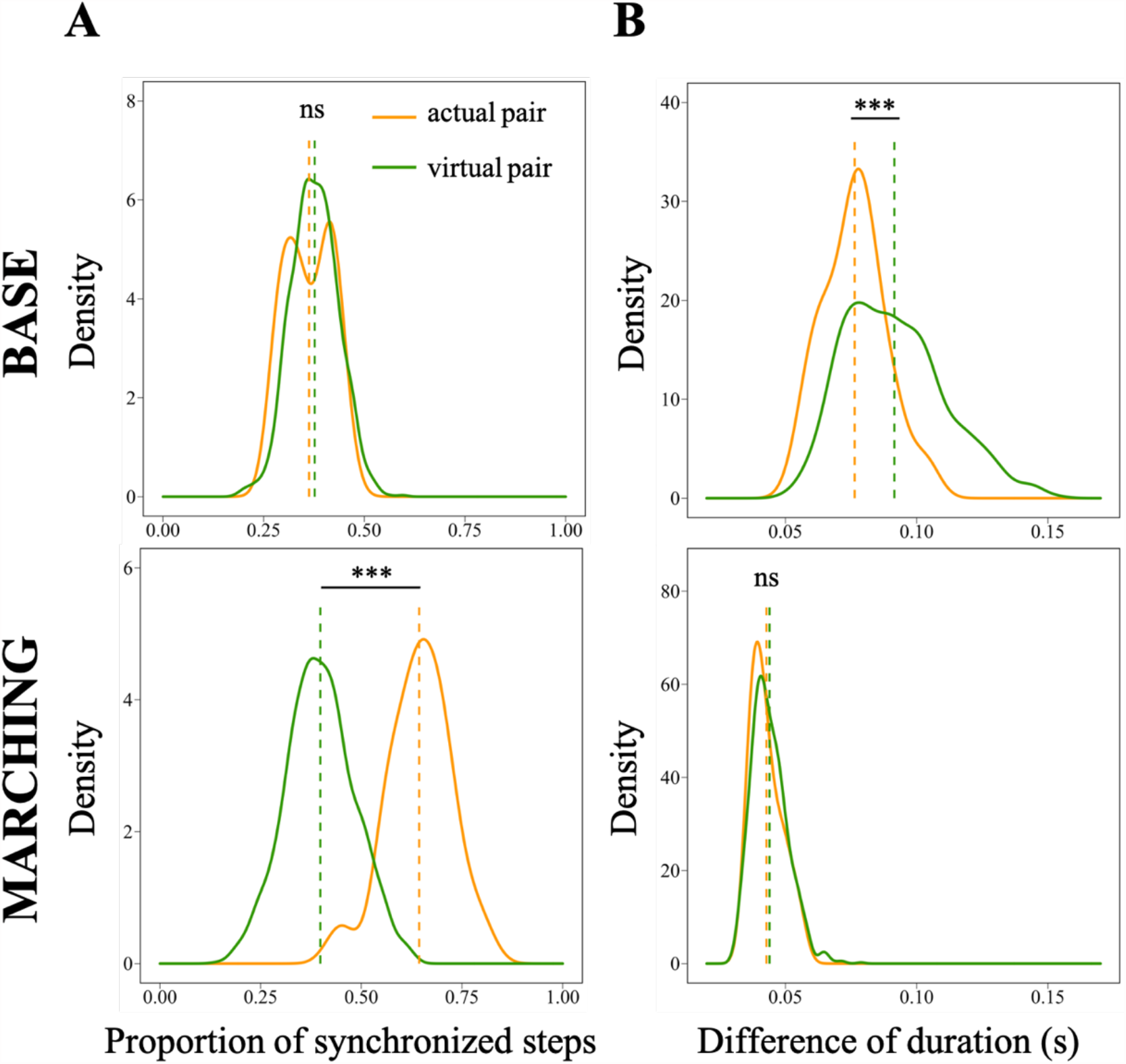
Footstep synchronization and duration coordination. Probability density functions of (A) proportion of synchronized footsteps and (B) differences of duration between two pedestrians in pairs under the (Top) BASE and (Bottom) MARCHING conditions. Orange (green) solid lines represent distributions of actual (random) pairs, respectively. Dashed vertical lines show the mean values of distributions. Asterisks indicate the statistical significance (\*\*\**P* < 0.001; ns, *P* > 0.05).

In general, when steps between two paired pedestrians are synchronized, their step durations (or frequencies) also tend to match, but the reverse does not always hold true; their step durations may match without synchronization (for example, in the case of a constant delay). Therefore, in addition to the synchronization, we also examined the degree of similarity in step duration between the two. We defined the step duration of a single foot for each pedestrian as the time between two consecutive heel strike timings. As an indicator of duration similarity, we simply determined the mean difference in step duration between the two paired pedestrians for each trial. The smaller the difference, the higher the similarity in step duration of pairs. The degree of similarity in duration in MARCHING was considerably smaller than that in BASE (Fig. 2B). Also, the degree of duration similarity of actual pairs was significantly smaller than that of random pairs in BASE (Welch’s *t*-test, *N*_*actual*_ = 20, *N*_*random*_ = 1000, *t* = −5.55, *p* < 0.001, Cohen’s *d* = −0.79), but there was no difference in MARCHING (Welch’s *t*-test, *N*_*actual*_ = 20, *N*_*random*_ = 1000, *t* = −0.88, *p* = 0.38, Cohen’s *d* = −0.17). This implies that pedestrians spontaneously coordinated their footstep durations to some extent in the BASE condition and pedestrians simply matched it via the shared external cues rather than as the result of coordination in the MARCHING condition.

To summarize, groups with the external auditory cue significantly synchronized their stepping motions, whereas groups without the external cue coordinated their durations but not their step synchronization.

### Influence of external auditory cues on self-organization of bidirectional flow

According to our observations, pedestrians under the BASE condition seemed to deviate laterally more and merge into fewer lanes (i.e., wider lanes) than those under the MARCHING condition. In this section, we quantitatively evaluate the width of lanes and lateral deviation and show their relation to the robustness of lane formation. To this end, we analyzed pedestrian behavior in more detail by dividing the time development of lane formation into five independent (non-overlapping) stages [5] from the beginning of the test through lane formation and dissolution as described in the Methods.

To evaluate the width of lanes, we calculated the average number of lanes in each trial by using a clustering method [4] based on the same follower–predecessor relationship used to define the pairs in the previous section. In short, if one pedestrian follows the other, those two pedestrians belong to the same cluster at a given moment of time (Methods). The number of lanes at a certain time is defined by the number of clusters in the central region of the measurement area (3 × 3 m, Fig. S2). We calculated the average number of clusters from stage 2 to stage 4 as the number of lanes for each trial and found that pedestrians under MARCHING organized significantly more lanes (i.e., thinner lanes) than those in BASE (Fig. 3, Welch’s *t*-test, *N*_*BASE*_ = 20, *N*_*MARCHING*_ = 20, *t* = −3.64, *p* < 0.001, Cohen’s *d* = −1.15).

**Fig. 3.**
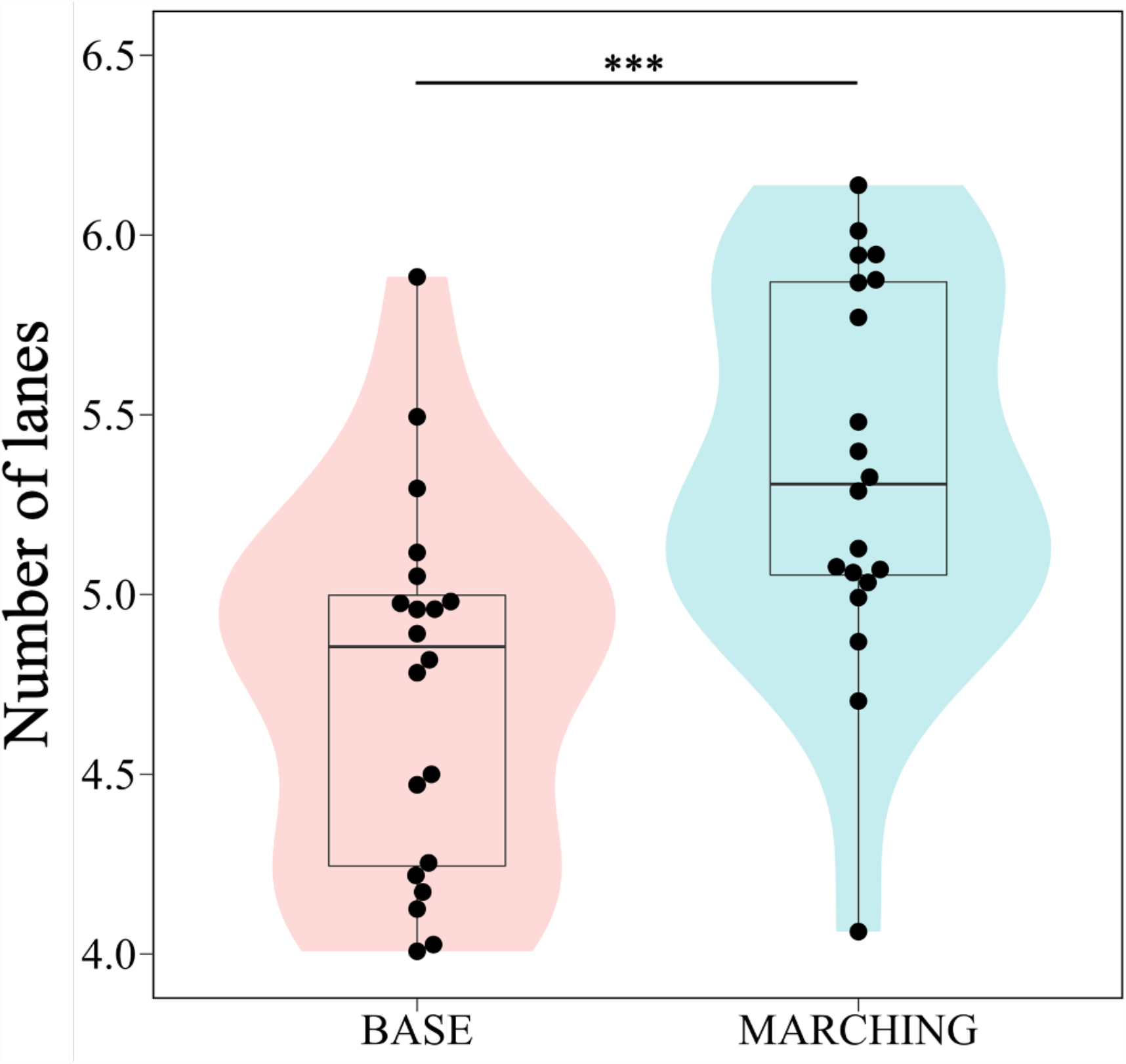
Mean number of generated lanes in each trial. Each datapoint represents a trial. Asterisks indicate the statistical significance (\*\*\**P* < 0.001). Box-and-whisker plots represent the median of the data (central thick line), the first and third quartiles (box), and 1.5× the interquartile range of the median (whiskers). The shaded areas show violin plots of the data.

We also investigated the potential collision risk of pedestrians to evaluate the intrinsic instability of lanes observed in our experiments because the width of lanes has been shown to directly influence the instability of lane structure [21]. To do so, we calculated the expected time to potential collision (τ), which is the duration of time for which two pedestrians in different groups could continue walking at their current velocities before colliding [22] (Methods). We also calculated the expected distance to potential collision, which is the distance traveled in *τ*. The smaller the time/distance to collision among pedestrians in a crowd, the more easily the ordered state would be disrupted by small perturbations. The results revealed that both time and distance to potential collision under MARCHING were significantly smaller than those under BASE (Fig. 4, Welch’s *t*-test, time to potential collision; *N*_*BASE*_ = 20, *N*_*MARCHING*_ = 20, *t* = 5.27, *p* < 0.001, Cohen’s *d* = 1.66, distance to potential collision; *N*_*BASE*_ = 20, *N*_*MARCHING*_ = 20, *t* = 4.45, *p* < 0.001, Cohen’s *d* = 1.41), suggesting that pedestrians in BASE could create a more robust structure than those in MARCHING.

**Fig. 4.**
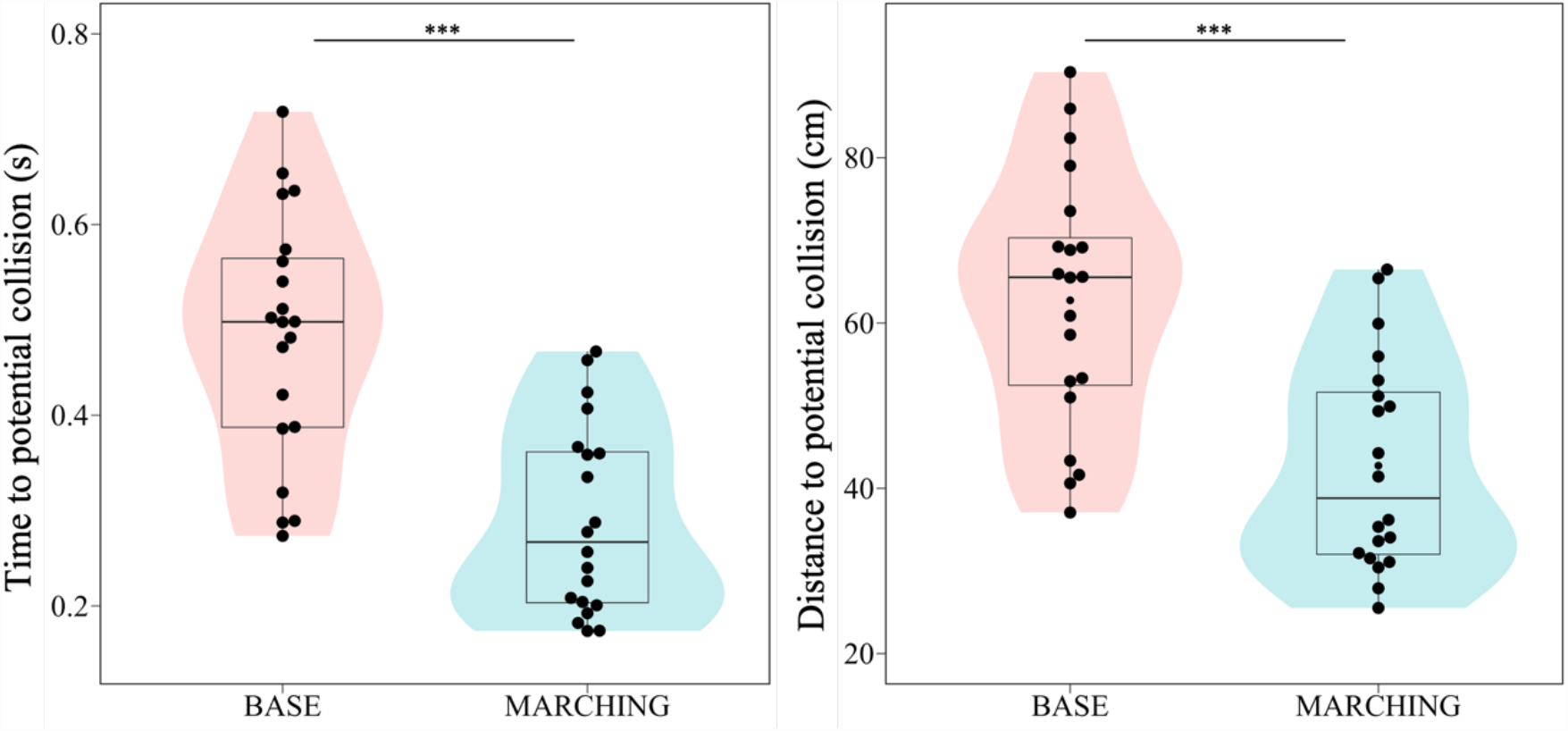
Mean time (left) and distance (right) to potential collision in each trial. Each datapoint represents a trial. Asterisks indicates the statistical significance (\*\*\**P* < 0.001). See Fig. 2 for a description of the box-and-whisker plots. The shaded areas show violin plots of the data.

Finally, we investigated the lateral fluctuations/deviations of pedestrians from the direct straight path to their destination. This deviation is well captured by the curvature (*κ*) of walking trajectories [23] (see Methods). A larger *κ* indicates a greater deviation, while *κ* = 0 indicates a completely straight trajectory. We observed that the mean curvature for each trial in BASE was significantly larger than that in MARCHING in all stages except stage 3 (Fig. 5, Welch’s *t*-test, stage 1: *t* = 2.19, *p* = 0.034, Cohen’s *d* = 0.69; stage 2: *t* = 3.63, *p* < 0.001, Cohen’s *d* = 1.15; stage 3: *t* = 0.83, *p* = 0.40, Cohen’s *d* = 0.26; stage 4: *t* = 2.80, *p* = 0.008, Cohen’s *d* = 0.88; stage 5: *t* = 3.79, *p* < 0.001, Cohen’s *d* = 1.20; *N*_*BASE*_ = 20 and *N*_*MARCHING*_ = 20 for each stage except for stage 3 under MARCHING where *N*_*MARCHING*_ = 19, see Methods). Importantly, in the middle of lane formation (i.e., stages 1 and 2), pedestrians in BASE deviated more than they did in the same stages in MARCHING, suggesting that they can self-organize into wider lanes the more they deviate from a straight-line trajectory. This suggests that the pedestrians who were forced to synchronize their steps had a more limited ability to perform their usual exploratory walking to form robust lanes.

**Fig. 5.**
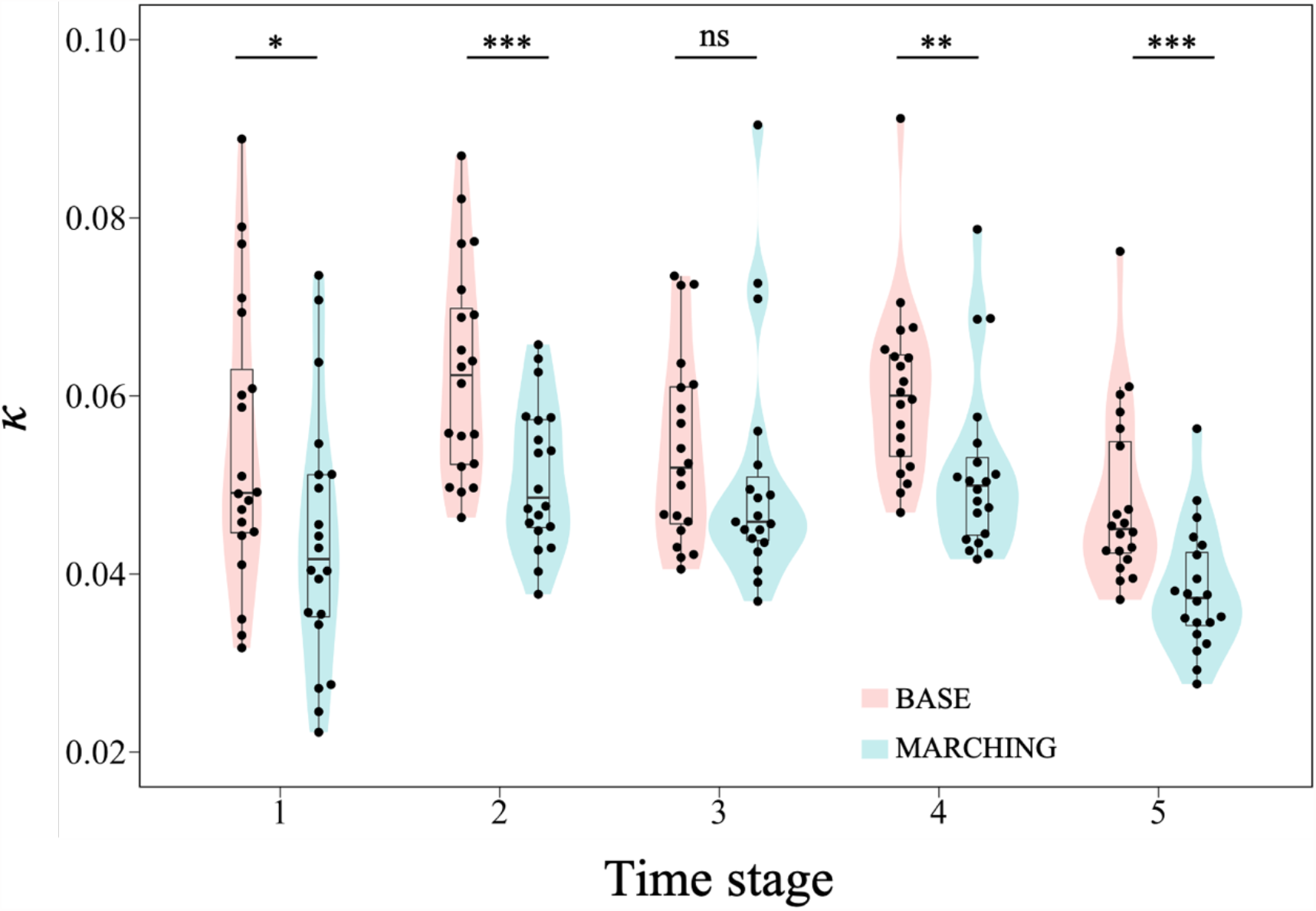
Mean curvature of pedestrians’ trajectories (*k*) during each stage of time development of lane formation. Each datapoint represents a trial. Asterisks indicates the statistical significance (\*\*\**P* < 0.001; \*\**P* < 0.01; \**P* < 0.05; ns, *P* > 0.05). See Fig. 2 for a description of the box-and-whisker plots. The shaded areas show violin plots of the data.

## Discussion

In this study, we addressed whether footstep synchronization occurs in pedestrian crowds with relatively more freedom of movement and contributes to spatial self-organization. To this end, we conducted bidirectional flow experiments. One condition had external auditory stimuli to be followed by pedestrians and the other did not. Footstep synchronization was only observed in the former condition. This synchronization driven by the external auditory cues increased the number of lanes formed, causing structural instability and potential collision risks. It also restricted the pedestrians’ usual exploratory lateral movements, especially in the middle of the lane formation process. These results imply that the pedestrians with no external conductors did not spontaneously synchronize; instead, they were able to deviate laterally, which appeared to promote robust formation of groups. In contrast, the footstep synchronization driven by the external cues disturbed the process of lane formation.

In previous studies, pedestrians have been shown to synchronize their foot motions in single-file experiments [12–14], where the mechanism for synchronization was attributed to pedestrians stepping in unison on the same line to efficiently exploit spatial resources and avoid collisions with preceding pedestrians [14]. However, a prerequisite for this mechanism to function (i.e., the single-file structure guaranteed by the boundary condition) was not met under our experimental conditions, which allowed for more freedom of movement. In our scenario, pedestrians can easily move laterally to avoid collisions. Moreover, overtaking behavior and the presence of oncoming pedestrians also lead to lateral movements, distracting pedestrians from solely focusing on preceding pedestrians. In addition, pairs of pedestrians in a non-crowded open space spontaneously synchronize their stepping, presumably through auditory feedback of steps [10, 11]. However, it would be difficult for pedestrians to hear the sound of steps in our crowded experimental conditions. Finally, it is possible that when pedestrians are moving toward each other, asynchronous stepping would facilitate smooth collision avoidance, because pedestrians who move later find it easier to adjust their movements relative to those who moved first [24]. Thus, it seems rather natural to consider that pedestrians in a crowd with more freedom of movement take steps asynchronously. Most footstep synchronization in crowds is likely to be induced by indirect interactions such as the pedestrians’ interactions observed in the crowd accident on Millennium Bridge [16].

Our study revealed that the temporal pattern of stepping motions influenced the spatial motions of pedestrians. At the individual level, we observed that the variability of lateral fluctuations of an individual’s trajectory from the direct path to the destination decreased when pedestrians followed the external auditory cue (Fig. 5). At the collective level, the external cue increased the number of generated lanes (Fig. 3). Lateral movements can play an important role in the process of pattern formation. In the condition with no cues, moving laterally enables pedestrians that are more distant from each other in the same group to come closer, which contributes to formation of wider lanes. In the condition with external auditory cues, there is less lateral movement, allowing pedestrians to cut between oncoming pedestrians if there is enough space, thereby creating more (thinner) lanes. Moreover, the analysis on time/distance to potential collisions (Fig. 4) supports the idea that the width of lanes corresponds to the robustness of generated structure. This result is consistent with a previous theoretical study showing that a larger number of lanes (i.e., narrower lanes) have a shorter lifetime because of the larger fraction of pedestrians at contact surfaces between lanes moving in the opposite direction [21]. Our findings therefore highlight the possibility that external cues disturb the process of lane formation and emphasize the importance of exploratory lateral movements in promoting robust self-organization in human crowds.

In addition, pedestrians spontaneously coordinated their steps in terms of duration (or frequency). Indeed, without external cues, the differences in duration between two pedestrians in actual pairs were significantly smaller than those in random pairs (Fig. 2B top). Although additional data are required, this coordination may occur because two paired pedestrians in the same group have to adjust their movements when merging into a single lane so that they are not cut off by oncoming pedestrians. On the other hand, with external cues, the differences in durations of actual pairs were relatively narrowly distributed, but there was no significant difference in the durations of actual and random pairs (Fig. 2B bottom). This simply indicates that pedestrians followed external cues well, and hence, step frequencies did not vary much for any individual in the crowd.

Human locomotion is basically updated by each discrete footstep due to the biomechanics of the bipedal gait [25]. This can be represented as a discrete positional update in computational models. Most previous models of human crowd behavior assume synchronous position updates, in which all pedestrians simultaneously update their positions. However, for other collective animal behaviors, various theoretical models with asynchronous positional updates have been proposed [26–32]. For example, asynchrony in position updates has been suggested to allow anisotropy to emerge in interactions among individuals and to generate inherent noise, which drives autonomous motion in groups without external noise, as has been observed in real animal groups [28, 29]. Because we observed asynchronous movements among pedestrians and their influence on self-organization in human crowds, it would also be valuable to incorporate characteristics of asynchronous behavior in computational models in pedestrian crowds. In some theoretical work on non-human animal collective behavior, asynchrony plays important roles for anticipatory interactions among individuals [30–32], and these are also fundamental in pedestrian interactions [7]. We expect that our results may provide quantitative support to asynchronous models that incorporate anticipation. Enhanced human crowd models that integrate asynchronous behaviors encompassing lateral movements will offer a more accurate depiction and prediction of pedestrian flows.

There are, however, limitations in this work. In our study, experiments were conducted on bidirectional flow to investigate the relation between footstep synchronization and emergent pattern formation. Therefore, we did not test whether synchronization occurs in the unidirectional flow of a single group. Moreover, while we adopted a previously used experimental setting for bidirectional flows [5–7], we did not verify the dependence of footstep synchronization with the density of crowds. Furthermore, our experiments focused on comparatively short-term bidirectional flow, similar to that observed at busy crossings. However, there can be long-term bidirectional flow, for example, on streets with many shops, where the structural instability of lanes manifests [4] and lateral movements potentially have more influence than short-term bidirectional flow.

## Materials and Methods

### Experimental design

The experiment was conducted in May 2022 in the gymnasium of the Kyoto Institute of Technology.

### Participants

Forty-eight university students were recruited for the study (35 males, 13 females, mean (±SD) age = 21.08 ± 1.71 years). For ease of video analysis, each participant was asked to wear a black T-shirt; ID stickers were attached on both the right and left shoulders, and each participant wore a colored (red or yellow) cap. Written informed consent was obtained from all participants prior to the start of the study. The study was approved by the Ethics Committee of Kyoto Institute of Technology.

### Apparatus

The experimental corridor design was adapted from that described in previous studies [5–7]. There was a straight corridor consisting of three main parts: a measurement area (10 × 3 m) in the center of the corridor, with a waiting area (8.4 × 3 m) on each side of the measurement area. There were also buffer zones (1.2 × 3 m) between the measurement and waiting areas to enable participants to reach a stable walking speed before entering the measurement area (Fig. S1). As starting lines in the experiments, eight lines were drawn perpendicular to the long side of the corridor in each waiting area at an equal distance of 1.2 m. Destination areas were set at the end (outside) of each waiting area. The floor of the corridor was covered with a PVC gym floor cover so that participants could walk in their own outdoor shoes. Two loudspeakers presenting auditory stimulus to the participants were placed at a height of 0.76 m on either side of the measurement area, one for presenting instructions and the other for auditory stimulus.

### Procedure

The experimental procedure was similar to that used in previous experiments [5–7]; however, we revised some details to investigate the effect of an external conductor by means of auditory stimulus on footstep synchronization and on self-organization of bidirectional flow. First, the participants were divided into two groups of 24, each with an approximately equal gender ratio. Then, participants in each group were randomly positioned onto one of the eight starting lines, with three participants per line. At the start signal, they were asked to start walking toward the opposite end of the corridor and to keep walking until entering the destination area on the opposite side.

In addition to observing the pedestrians’ step synchronization during self-organization of lane formation, we attempted to observe the influence of instructions to follow external auditory stimuli. We therefore set two experimental conditions: baseline (BASE) and MARCHING. In the MARCHING condition, participants were asked to walk following the rhythm from an electric metronome (120 beats/min, approximately equal to the normal pedestrian footstep frequency [19]) played from the loudspeaker. In the BASE condition, they walked without the sound stimulus. To observe foot synchronization, an inertial measurement unit (IMU, AMWS020, ATR-Promotions, 100Hz) was attached just above the ankle to each leg of all participants in one of the two groups. After the IMUs were attached, the participants were asked to walk a round-trip (one by one) in the measurement area to acclimatize them to the area and synchronize the timestamps between the overhead camera and the other pedestrians’ IMUs (see the synchronization section for more details).

Before conducting the main experiments, two pretest trials were conducted, one for each condition, to confirm participants had correctly understood the instructions and could clearly listen to the auditory stimulus while they walked. The main experiments were divided into two blocks. As mentioned previously, we divided the participants into two groups, referred to as Group A and Group B. We asked all participants in Group A to attach the IMUs and they continued to wear them during the first experimental block. Then, the IMUs were removed from the participants in Group A and were attached to all participants in Group B for the next block. In each block, 20 trials were conducted (10 for each condition) in a randomized order, so experiments were replicated 20 times under each condition.

### Video tracking

The experiments were recorded from above with GoPro Hero 10 camcorders equipped with a linear digital lens (4K, 30fps) fixed at a height of 10 m. From the video images, we obtained time series pedestrian trajectory data using Petrack [33]. Participant IMU IDs in the trajectory data were confirmed by comparing them with the shoulder ID stickers.

### Synchronization of the timestamps between the body and foot movement tracking systems

To synchronize timestamps between the overhead video for body movement tracking and the IMUs for foot movement tracking, we annotated a time instant of each participant’s heel strike during the one-by-one test trip both on a frame of the overhead camera and a timestamp of the IMU using ELAN annotation software [34]. We annotated three different heel strikes for each participant to check for annotation errors. For IMU data, a heel strike was defined as the time at which the *Y* component of acceleration takes the local maximum value [35]. Timestamps of all IMUs were then adjusted to correspond with those of the overhead camera.

### Estimation of the timing of heel strike

To investigate footstep synchronization between a pair of two pedestrians, we measured the participants’ foot motions by means of the IMU attached to the participants. The *X, Y*, and *Z* axes of the IMUs were set to the craniocaudal, anterior–posterior, and medial–lateral directions, respectively. We used the timing of the heel strike to evaluate footstep synchronization and used an angular velocity in the medial–lateral direction to estimate it, as proposed in [36, 37]. To do this, we first synchronized the timestamps between all IMUs and the overhead video camera as described in the previous section. A median filter with a window length of 5 was then applied to the time series of the IMU data. Then, the timings of the heel strikes were extracted by searching for the local maximum sandwiched between the toe off and heel strike (for details, see [36, 37]). Local maxima were detected by using the SciPy *signal*.*find_peaks* function (version 1.9.1).

### Definition of a pair of pedestrians

We considered that pedestrian *i* follows pedestrian *j* at time *t* if the position of *i* at *t* is less than *dr* from the position of *j* at *t* – *dt*, with *dt* =1 s and *dr* = 0.7 m [4]. We then assumed that *i* and *j* are a pair if *j* was the individual followed by *i* for the longest amount of time during a trial. The first step to the last step of the preceding pedestrian in the measurement area was taken into account. The preceding pedestrian’s single (right or left) foot was paired with the corresponding follower’s one that had the nearest heel strike timing.

### Definition of footstep synchronization

As in previous studies [12–14], we employed the conformity characteristics of the timing of heel strikes (i.e., |*t*_*f*_ − *t*_*p*_| ≤ *K*) to define the synchronization of a single foot of a pair, where *t*_*f*_ and *t*_*p*_ are the timings of the heel strike of a single foot of the follower and the predecessor, respectively, and *K* is a short relaxation time about the difference of the heel strikes and is set to 0.1 s [10]. In addition to in-phase synchronization, we also considered anti-phase synchronization (i.e., |*t*_*f*_ – *t*_*p*_ + (*Δt*_*f*_ + *Δt*_*p*_)/4| ≤ *K* and |*t*_*f*_ − *t*_*p*_ − (*Δt*_*f*_ + *Δt*_*p*_)/4| ≤ *K*), where *Δt*_*f*_ and *Δt*_*p*_ are durations of the step of a single foot of the follower and the predecessor, respectively. We also took into account the conformity characteristics of duration (i.e., |*Δt*_*f*_ – *Δt*_*p*_| ≤ *K*_*d*_), where *K*_*d*_ is a short relaxation time about the difference of the durations and is set to (*Δt*_*f*_ + *Δt*_*p*_)/8 [12–14]. We then calculated the proportion of in-phase and anti-phase synchronized footsteps satisfying the duration conformity out of all footsteps for each trial as a degree of synchronization. Note that in our experiments, almost all footsteps hold the characteristics of this duration conformity.

### Random shuffling of pedestrian pairs

We generated 1000 virtual trials for each condition, where two pedestrians in pairs were randomly shuffled (i.e., a random pairing of the follower from one pair and the predecessor from another pair in the same condition), and analyzed the degree of synchronization of these random-pair trials in comparison with that of the original dataset. To generate virtual trials, for each pair in each trial, the pedestrian walking behind was fixed and the preceding pedestrian was randomly shuffled as follows: (i) one trial was randomly selected from all trials in the same condition except the focal trial (i.e., from 19 trials), and (ii) one pair was randomly selected from all pairs in this randomly selected trial except pairs containing members of the original pair, and the preceding pedestrian in this randomly selected pair was coupled with the original following pedestrian as a random pair.

### Definition of stages for time development of lane formation

We used the stages leading to the formation of lanes that were first proposed in [5]. To do so, we first employed six different time instants: *t*_0_: a pedestrian enters the measurement area for the first time; *t*_1_: the lead pedestrians of both groups crossed each other; *t*_2_: a pedestrian leaves the measurement area for the first time; *t*_3_: the last pedestrian enters the measurement area; *t*_4_: the last pedestrians of both groups crossed each other; and *t*_5_: the last pedestrian leaves the measurement area. Each interval between consecutive two time instants is considered to represent a different stage of lane formation: stage 1 (*t*_0_ – *t*_1_), two separate unidirectional flows until they intersect; stage 2 (*t*_1_ – *t*_2_), lane development in which pedestrians in the two groups start to separate into lanes; stage 3 (*t*_2_ – *t*_3_), full bidirectional flow, where the whole measurement area is filled with pedestrians of both groups; stage 4 (*t*_3_ – *t*_4_), lane dissolution in which lanes are being dissolved and there is unidirectional flow in parts of the measurement area; and stage 5 (*t*_4_ – *t*_5_), two separate unidirectional flows have re-emerged. Note that in one MARCHING trial, stage 3 was not defined because *t*_2_ took a larger value than *t*_3_, and hence we obtained 19 samples of stage 3 for MARCHING and 20 samples for the others.

### Curvature of pedestrian’s walking trajectory

To evaluate pedestrian’s deviation from the direct straight path to the destination, we calculated time series of curvatures of trajectories [23]. Curvature *κ* at time *t* was defined as a norm of the difference between unit vectors of velocity at time *t* and *t* + *dt*,

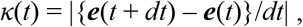

where ***e***(*t*) = ***v***(*t*)/|***v***(*t*)|, ***v***(*t*) = ***r***(*t*) – ***r***(*t – dt*), and ***r***(*t*) is a position of an individual at time *t. dt* was set at 1 s [23]. A larger *κ* indicates a greater pedestrian trajectory deviation, whereas *κ* = 0 indicates a completely straight trajectory. The mean curvature of each stage in each trial was calculated and used as an indicator of the degree of lateral deviation in pedestrian trajectories.

### Potential collision risk

We defined potential collision risk in time as the time to collision (denoted as *τ*), which was calculated as the duration of time two pedestrians could continue walking at their current velocities before colliding. Mathematically, for any two potentially colliding pedestrians *i* and *j*,

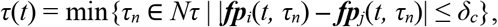

where ***fp***_*i*_(*t, τ*_*n*_) = ***r***_*i*_(*t*) + *τ*_*n*_ · ***v***_*i*_(t), *Nτ* = {0.03, 0.06, …, 9.0}, and *δ*_*c*_ is the collision detection threshold (set to 0.6 m). Potential collision risk in space (distance to collision) was then calculated as *λ*(*t*) = *τ*(*t*)·{|***v***_*i*_(*t*)| + |***v***_*j*_(*t*)|}/2.

### Statistical analyses

Welch’s *t*-test was used to compare means for relevant analyses between conditions and actual and random datasets using R version 4.0.3 (The R Foundation for Statistical Computing, Vienna, Austria).

