## Supplemental Figures for "Robust spatial self-organization in crowds of asynchronous pedestrians"

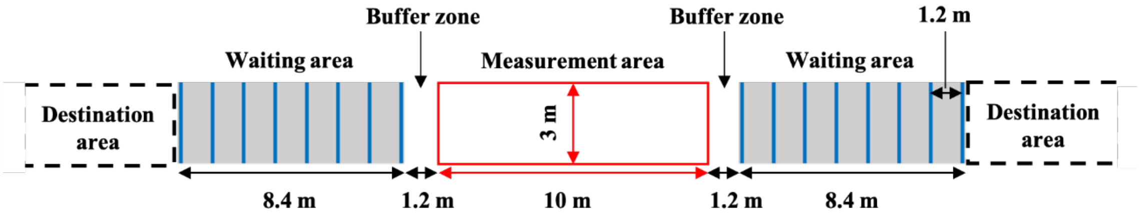


**Fig. S1. Illustration of the experimental corridor.** There was a straight corridor consisting of three main parts: a measurement area (outlined in red) in the center of the corridor, with a waiting area (gray) on each side of the measurement area. There were also buffer zones between the measurement and waiting areas to enable participants to reach a stable walking speed before entering the measurement area. Each waiting area includes eight lines used as starting locations (vertical blue lines). Destination areas were set at each end, outside of the waiting areas.

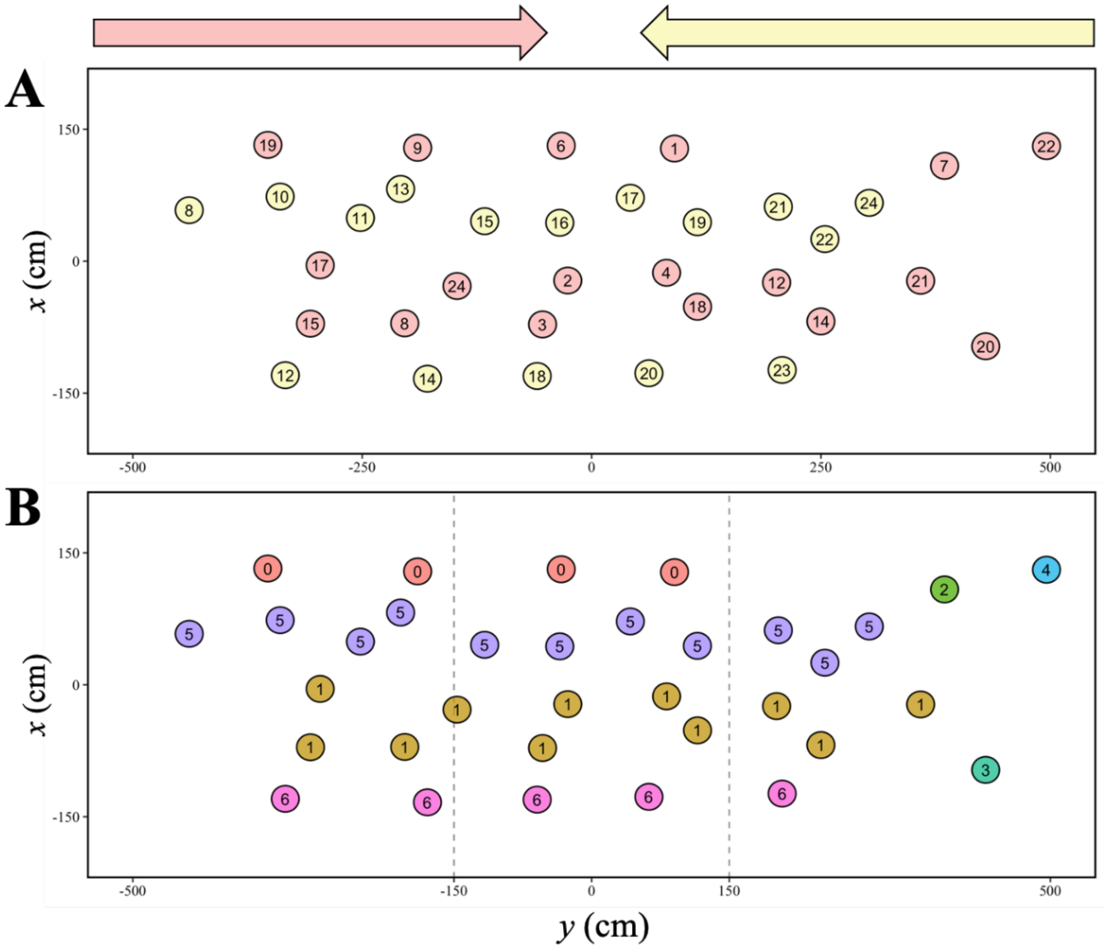


**Fig. S2. Illustration of the clustering method.** (A) Reconstructed pedestrian positions of two groups at a given time. Yellow (red) circles represent pedestrians moving from right (left) to left (right). The number in a circle represents the individual's ID. (B) Results of the clustering methods. Different colors and numbers in a circle correspond to different clusters. The vertical dashed lines indicate the central region of the measurement area. We assumed that the number of lanes is the number of clusters present in the central region. In this snapshot, the number of lanes was four.
